# Scavenger receptor CD163 multimerises to allow uptake of diverse ligands

**DOI:** 10.1101/2024.12.18.629151

**Authors:** Richard X. Zhou, Matthew K. Higgins

## Abstract

Scavenger receptors mediate cellular uptake of diverse ligands. CD163 is an archetypal scavenger receptor with a primary role in haemoglobin detoxification. Haemoglobin released from lysed erythrocytes causes oxidative damage. This is countered by formation of haemoglobin-haptoglobin complexes and CD163-mediated uptake into macrophages for degradation. Different haptoglobin isoforms form stoichiometrically diverse haptoglobin-haemoglobin complexes which must all be recognised by CD163. We show that CD163 forms multimeric complexes, with calcium-mediated interactions forming a trimeric base. Arms emerge from this base, moulding around the ligand to form a single promiscuous ligand-binding site. The arms can also bind to arms of adjacent protomers, blocking the ligand-binding surfaces in an autoinhibited state. This results in a scavenger receptor that can recognise diverse haemoglobin-containing complexes, allowing physiologically important detoxification.

## Introduction

Transport of oxygen in mammalian blood is mediated by haemoglobin (Hb) tetramers which circulate within erythrocytes. But free Hb is toxic, due to the reactive properties of the haem group, which can engage in oxidative reactions and generate free radicals (*1-3*). Hb can be released from erythrocytes due to haemolysis or tissue damage and free Hb increases in chronic conditions such as sickle cell anaemia (*4*) or during infections such as malaria (*5*). Mammals have therefore evolved a system to detoxify free Hb.

Hb detoxification involves the abundant serum protein, haptoglobin (Hp). The serine protease (SP) domain of Hp binds to an Hb αβ-dimer (*6, 7*). Hp also contains complement control protein (CCP) domains, which differ in number in the two isoforms found in humans (*8*). In people homozygous for isoform 1, a single CCP domain per Hp chain mediates dimer formation, creating a ‘dumbbell’-shaped haptoglobin-haemoglobin (HpHb) complex with two ‘heads’. In contrast, isoform 2 allows formation of higher-order oligomeric states in individuals who are either homozygous for isoform 2 or heterozygous (*9-11*). The formation of HpHb complexes buries reactive groups on Hb and reduces their capacity to cause oxidative damage (*6*).

The second step in detoxification is uptake of HpHb into macrophages. This is mediated by CD163 (*12*), which is the archetypal class I scavenger receptor. CD163 contains an ectodomain consisting of nine scavenger receptor cysteine-rich (SRCR) domains linked to a C-terminal transmembrane helix (*13*). CD163 must bind to very different architectures of HpHb complexes formed from different Hp isoforms and mediate their endocytosis (*10, 12*). In contrast, free Hp is not recognised (*12, 14*), ensuring that it is not depleted unnecessarily from the serum. Finally, HpHb complexes should be released in the endosomes (*12, 15*), allowing degradation and detoxification of Hb. CD163 is also exploited by viruses for host cell entry. These include the simian haemorrhagic fever virus, which has zoonotic potential (*16*), and the major livestock pathogen, porcine reproductive and respiratory syndrome virus (*17*), which is responsible for estimated annual losses of $600 million (*18*). While the structure of isoform 1 HpHb complex has been determined (*6, 19*), little was known about how CD163 selectively binds to a structurally diverse range of HpHb complexes, why it does not bind to Hp and how it releases its ligand. We therefore combined cryogenic electron microscopy, biophysics and cell-based uptake assays to understand the molecular mechanism of CD163-mediated ligand uptake and release.

## Results

### The structural basis for binding of haptoglobin-haemoglobin to CD163

To understand how human CD163 binds to HpHb, we expressed the complete ectodomain in HEK293 cells. Hp(1-1)Hb complexes were assembled by mixing human Hp isoform 1 with human Hb and were combined with CD163 ectodomain (Fig. S1a). The complex was purified by size-exclusion chromatography and grids were prepared for cryogenic electron microscopy. Data were collected on a Titan Krios and processed using SIMPLE (*20*) and CryoSPARC (*21*) (Fig. S2 and Table S1), resulting in two predominant three-dimensional classes. In both cases, we observed a single SP domain of Hp bound to an Hb αβ-dimer and this was coordinated by either two or three CD163 ectodomains (Fig. 1a,b). Indeed, size-exclusion chromatography with multi-angle laser light scattering (SEC-MALLS) showed concentration-dependent multimer formation in solution, with addition of HpHb increasing the fraction of multimeric complexes (Fig. S1b,c).

**Fig. 1.**
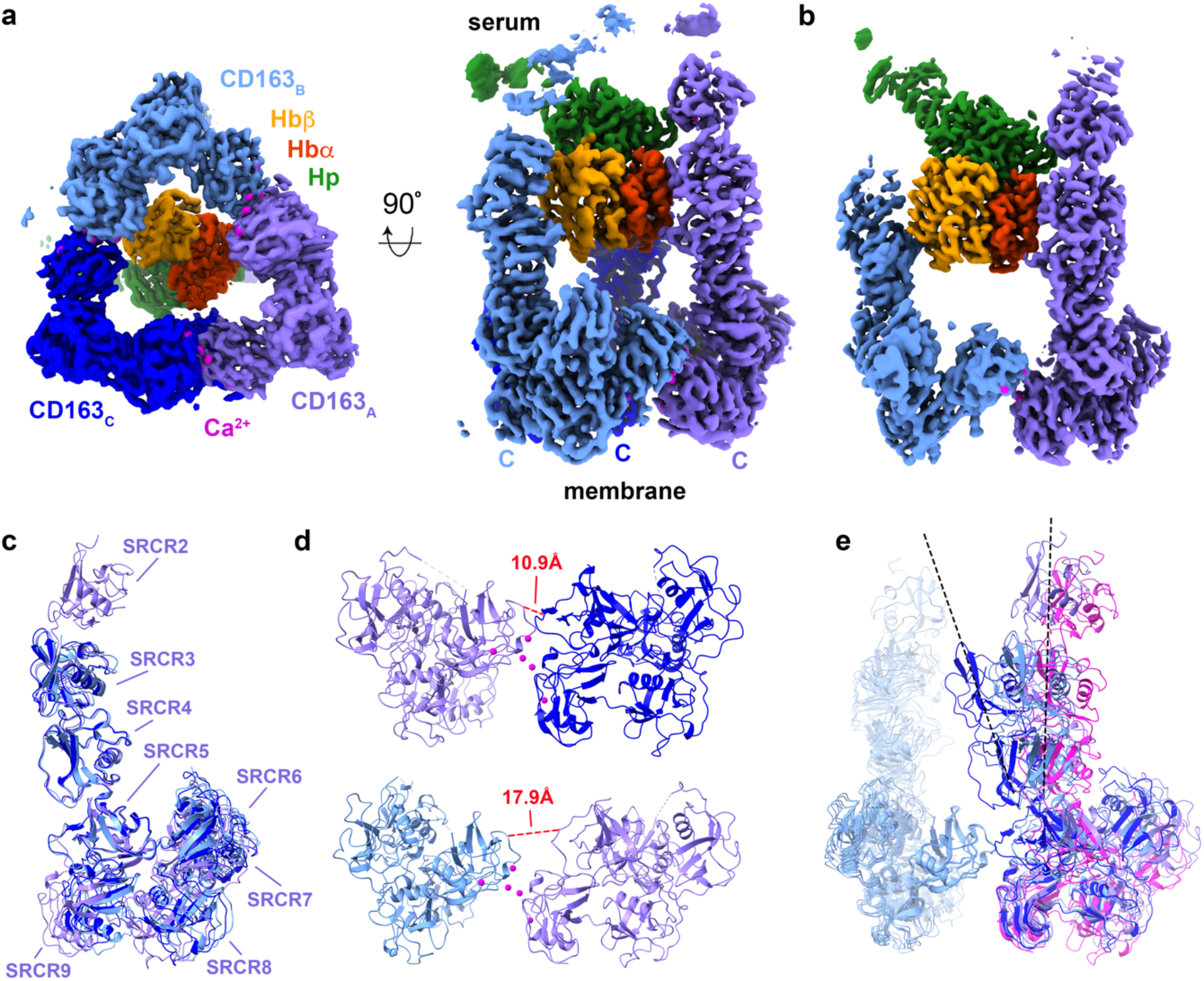
The structure of CD163 bound to HpHb. **a**. The structure of a trimer of the CD163 ectodomain bound to the HpHb complex. The three copies of CD163 are shown in three shades of blue. HpSP domain is green, the α-subunit of Hb is red and the β-subunit of Hb is orange. **b**. The structure of a dimer of the CD163 ectodomain bound to the HpHb complex, coloured as in a. **c**. An alignment of the three CD163 molecules found in the trimeric complex, aligned on SRCR5, with the SRCR domains labelled. **d**. A close-up of the interfaces between two pairs of CD163 subunits with the red dotted line showing the different distances between the Cα atoms of residues 561 and 804 in these two interfaces. **e**. An alignment of each pair of CD163 ectodomains observed in the trimeric and dimeric complex, with each aligned on SRCR7 of the left-hand CD163. The right-hand CD163 molecules from the trimer are coloured as in a. and in pink for the CD163_A_ from the dimer. Dashed lines indicate the maximum degree of tilting of the arm of CD163 in these structures.

The CD163 ectodomain adopts an architecture consisting of a ‘base’, formed from SRCR domains 5-9, linked to a four-domain ‘arm’ (SRCR1-4), of which two or three domains were ordered (Fig. 1c). The five domains of the base form a compact assembly which folds to bring SRCR5 and SRCR9 from the same monomer together. Multimer-formation is mediated by the base, predominantly due to interactions between SRCR7 of one subunit and SRCR9 of its neighbour (Fig. 1d). These interfaces contain spheres of density, which we attribute to be calcium ions, as calcium is required for HpHb binding and uptake (*12, 22*). The three interfaces within the trimer differ, with ‘rocking’ motions around these calcium ions leading to different relative positions of the two protomers (Fig. 1d).

The CD163 trimer has a flat triangular base, with the C-termini, which link to transmembrane helices, emerging from the flat surface (Fig. 1a). Protruding in the opposite direction are the arms, which extend away from the membrane. These three arms collectively form a binding site for one head of HpHb, with each arm interacting with a different surface of the head. The dimer adopts a similar global architecture, with the CD163_A_ arm making the same contacts with HpHb as in the trimer, but with the second arm making different interactions. While each arm adopts a very similar conformation (Fig. 1c), flexibility in their relative positions occurs due to the rocking motion within the base (Fig. 1e). This allows the arms to emerge from the base at different angles and to make different contacts with the asymmetric HpHb head. Flexibility of the CD163 base therefore allows its arms to mould to the ligand, likely allowing a scavenger receptor to create different binding sites for differently structured ligands.

### Structures of unliganded CD163 suggest a mechanism of autoinhibition

To determine whether the HpHb binding site is pre-formed in the absence of ligand, we also used cryogenic electron microscopy to reveal the structure of unliganded CD163. In this case, particles were distributed into three distinct three-dimensional classes (Fig. S3). The first class showed a trimeric arrangement of CD163 (Fig. 2a). Here, the base was well-resolved, adopting a similar conformation to that observed in the presence of ligand. In contrast, the density for the arms was poorly defined.

**Fig. 2.**
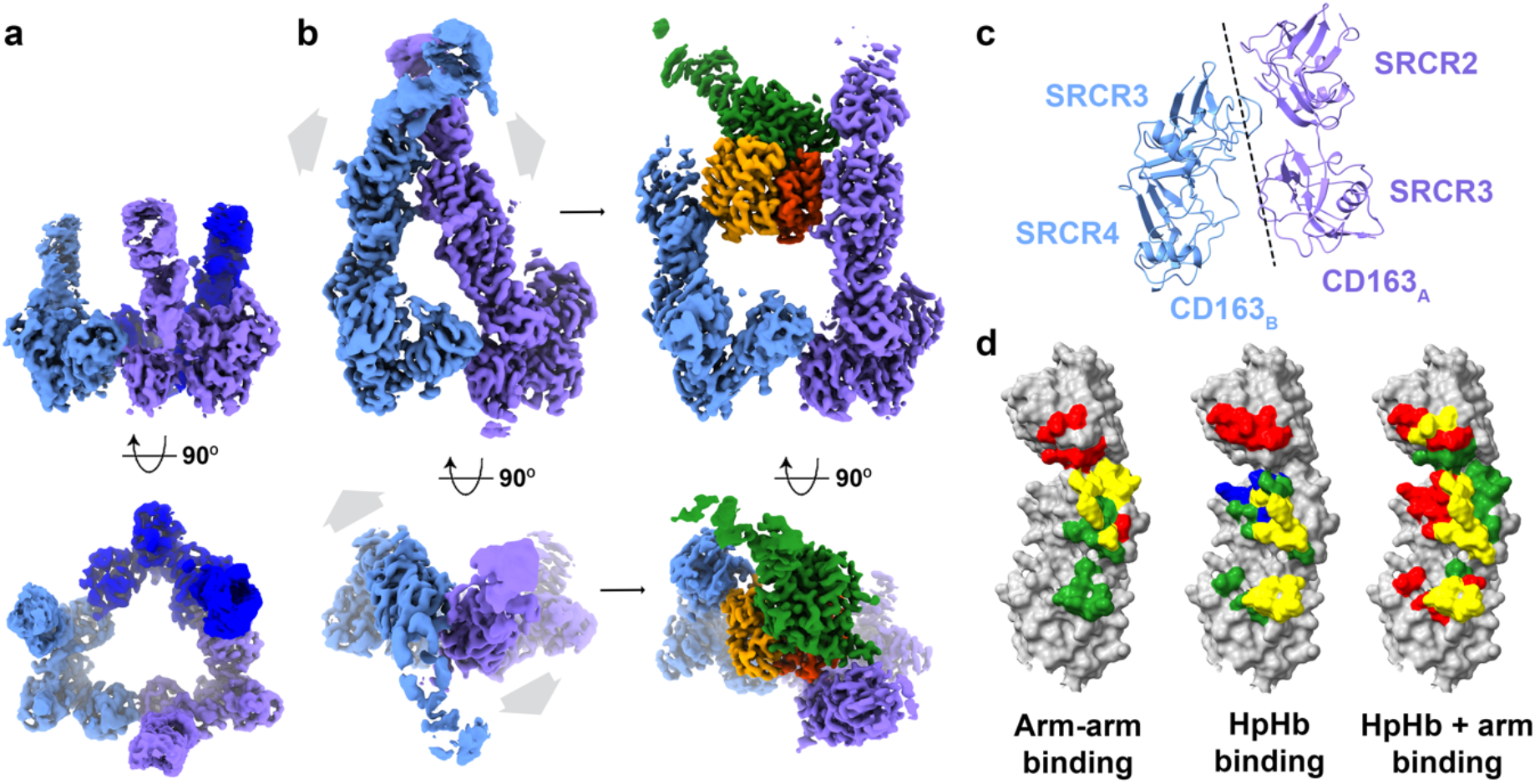
The structure of unliganded CD163. **a**. The structure of a trimer of CD163 in the absence of ligand, with disordered arms, coloured as in Fig. 1. **b**. The structure of a dimer of CD163 in the absence of ligand (left) and in the presence of HpHb (right), showing arm-arm contacts between the CD163 arms in the unliganded dimer, with the arms moving outwards to allow ligand binding. **c**. Close-up on the interface between the arms of CD163_A_ and CD163_B_ in the unliganded dimer. **d**. A schematic presenting the overlap in residues involved in arm-arm contacts and HpHb binding. The left panel shows SRCR2-4 of CD163 as a surface representation with residues involved in arm-arm contacts in the unliganded dimer coloured red for CD163_A_, green for CD163_B_ and yellow if involved in both subunits. In the central panel, residues are labelled if directly contacting HpHb in the liganded dimer and trimer. Red represents CD163_A_, green CD163_B_, blue CD163_C_, with yellow depicting residues involved in ligand binding in at least two CD163 subunits. The right-hand panel shows the same surface with residues labelled red if they contact HpHb, green if they are involved in arm-arm contacts in the unliganded dimer and yellow if both.

The other two three-dimensional classes revealed dimeric and trimeric arrangements of the unliganded receptor in which the arms of neighbouring protomers form contacts with each other. In the dimeric form, SRCR2 and SRCR3 of CD163_A_ interact with SRCR3 and SRCR4 of CD163_B_ (Fig. 2b,c). In the trimeric form, the same contacts between CD163_A_ and CD163_B_ are observed, with additional weaker density for the arm of CD163_C_, showing it to contact the arms of the other two subunits (Fig. S5). The surfaces of the SRCR domains which mediate contacts between the arms overlap with those which interact with HpHb (Fig. 2d). Contacts between arms will compete with ligand-binding interactions and the arms must move apart to create a ligand-binding site.

Unliganded CD163 on the surface of macrophages is therefore likely to exist in an equilibrium in which the arms are either available for ligand binding or interact together, occluding ligand binding. Interactions between arm and ligand will be in competition with interactions between neighbouring arms, providing a mechanism of autoinhibition. This could ensure that very weak ligands, which cannot compete with these arm-arm interactions, are not internalised.

### CD163 mediates uptake of ligands with different stoichiometries and structures

Analysis of the HpHb-bound trimer shows that each arm of CD163 interacts differently with HpHb (Fig. 3a). CD163_A_ interacts with the Hb α-subunit through SRCR3 and SRCR4, and with Hp through SRCR2. CD163_B_ recognises the β-subunit of Hb through SRCR3 and SRCR4 and CD163_C_ binds to the α-subunit of Hb through SRCR3 and SRCR4 and to Hp through SRCR3. Remarkably, it is the same faces of these SRCR domains which mediate different interactions with different surface features of HpHb, with the plasticity of the HpHb-binding site forming an asymmetric binding site, despite being formed from a homotrimer of receptors.

**Fig. 3.**
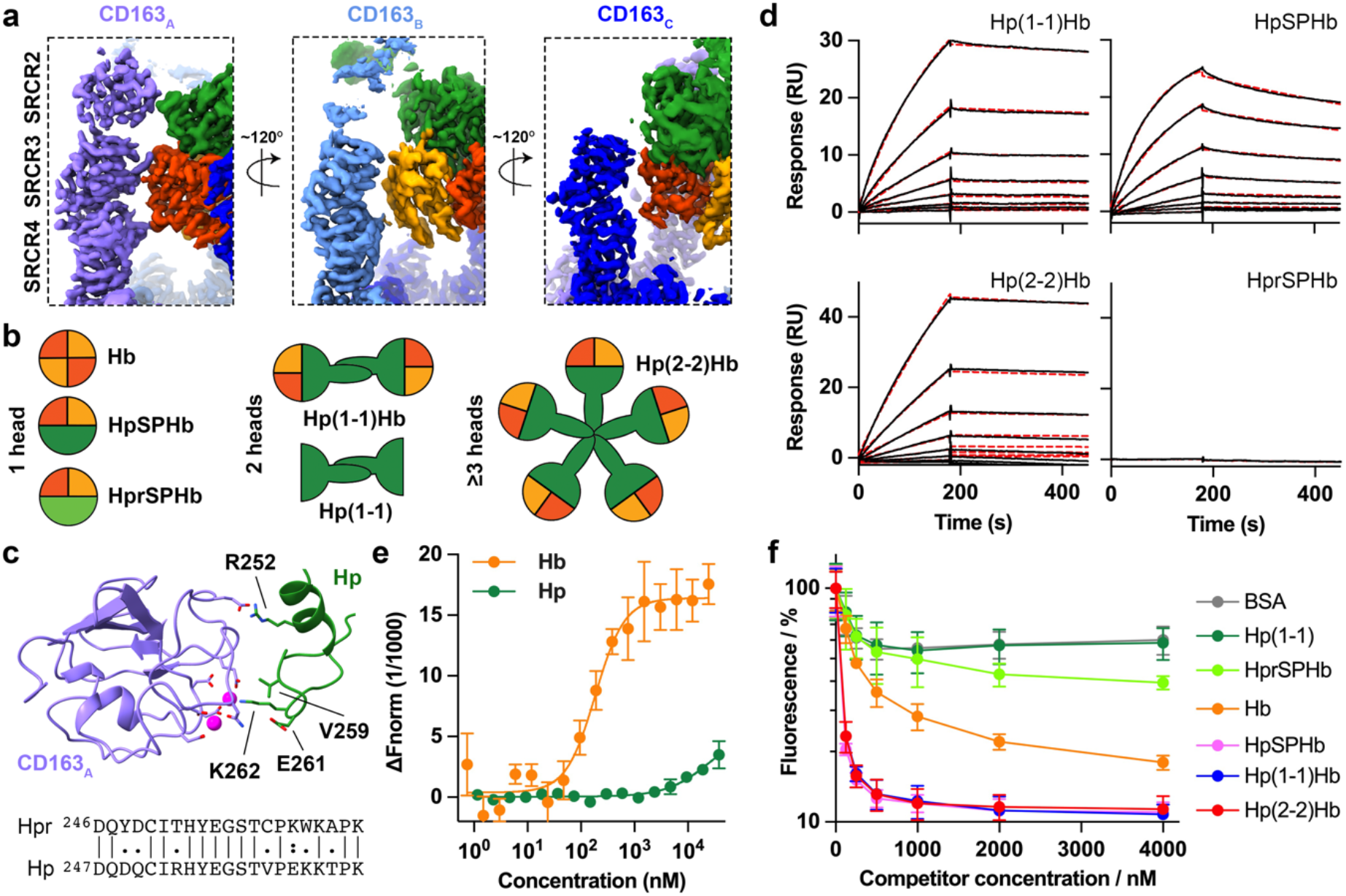
Ligand selectivity of CD163. **a**. The structure of the CD163 trimer bound to HpHb viewed from three directions, showing that each arm of CD163 interacts differently with the HpHb ligand. **b**. Schematic showing the different stoichiometries and structures of the ligands studied here. **c**. A view of SRCR2 of CD163_A_ (violet) bound to Hp (green). Below this is shown a partial sequence alignment of Hp and Hpr. Residues which differ between Hp and Hpr and which might influence binding to CD163 are shown as sticks and labelled. **d**. Assessment of the binding of CD163 ectodomain, immobilised through a C-terminal biotin, to Hp(1-1)Hb, Hp(2-2)Hb, HpSPHb and HprSPHb by SPR analysis. These represent two-fold dilution series from a maximum concentration of 10 nM for Hp(1-1)Hb and Hp(2-2)Hb, 20 nM for HpSPHb as well as an injection at 20 nM for HprSPHb. Data are shown as black lines while fitting to a one-to-one binding model is depicted as dashed red lines. These are representative of n = 3. **e**. Assessment of the binding of Hp and Hb to CD163 ectodomain using MST. Each point is the mean of three replicates and the error bars are the standard deviation. **f**. Measurement of the ability of different ligands to compete for the uptake of fluorescently labelled Hp(2-2)Hb into HEK293 cells transfected with CD163. BSA is included as a control for CD163-independent effects. Each point represents the mean of three replicates and the error bars are standard deviations.

HpHb complexes are found in a variety of multimeric states. The dumbbell-shaped Hp(1-1)Hb is formed when an Hp(1-1) dimer, presenting two protomers of HpSP, binds to an Hb αβ-dimer on each end (*6*). In contrast, higher-order multimeric states of Hp are found in people homozygous for isoform 2, leading to formation of Hp(2-2)Hb (*11*) (Fig. 3b). We observed that the three arms of CD163 come together to interact with a single head of HpHb, with weak density for the CCP domain of Hp projecting away from the membrane towards the other head (Fig. 1b). This leads to the hypothesis that HpHb complexes formed from different isoforms, and an artificial version containing just the HpSP domain bound to the Hb dimer (HpSPHb), would show similar binding and uptake properties. The structure also showed that the ∼68 % of the CD163 interaction surface on the HpHb complex is mediated by Hb subunits, with ∼32 % due to Hp, suggesting that CD163 might mediate Hb uptake more efficiently than that of Hp. Additionally, human sera contains haptoglobin-related protein (Hpr) (*23*) and a complex of Hpr and Hb is part of the trypanolytic factor which kills African trypanosomes (*19, 24, 25*). It is important that this is not taken up into human macrophages (*24*), where it might cause cell death. Polymorphisms between Hp and Hpr are found in residues which contact CD163 (Fig. 3c), suggesting that they might prevent binding. To test these predictions, we developed binding and uptake assays and assessed the outcomes for different ligands.

We first assembled Hp(1-1)Hb, Hp(2-2)Hb, HpSPHb and a complex of the SP domain of Hpr bound to Hb (HprSPHb). We coupled the CD163 ectodomain with a biotinylated BAP tag at the C-terminus to a surface plasmon resonance (SPR) chip surface, ensuring presentation with an orientation which matches that on a membrane. We then flowed the different HpHb complexes over the surface. Hp(1-1)Hb, Hp(2-2)Hb and HpSPHb bound with apparent affinities of 0.28, 0.16 and 1.3 nM, supporting the hypothesis that CD163 trimers recognise a single HpHb head (Fig. 3d, Fig. S6a and Table S3). In contrast, HprSPHb showed no binding at a concentration of 20 nM. This assay could not be used to measure the affinity for Hb and Hp, due to irregular shape of the SPR sensorgrams and to high non-specific binding. We instead used microscale thermophoresis (MST), which revealed affinities of 0.19 μM for Hb and 27 μM for Hp (Fig. 3e, Fig. S7 and Table S3). Therefore, HpHb binds to CD163 most tightly, followed by Hb, with little binding observed for Hp or HprSPHb.

In addition, we used a cell-based uptake assay in which HEK293 cells were transfected with full-length CD163 modified with a cytoplasmic GFP tag (*15*). We showed that uptake of fluorescently labelled Hp(2-2)Hb in HEK293 cells increases nearly 10-fold in the presence of CD163 (Fig. S8a). We then assessed the ability of different concentrations of heads of unlabelled ligands to compete for the uptake of 50 nM labelled Hp(2-2)Hb, as quantified by fluorescence-activated cell sorting (FACS) (Fig. 3f). Our BSA negative control and unconjugated Hp isoform 1 both showed a similar concentration dependence, suggesting no receptor-mediated uptake of Hp at 4 μM concentration. As Hp is normally found in sera at concentrations of 5-30 μM (*26*), receptor-mediated internalisation will be low. In contrast Hp(1-1)Hb, Hp(2-2)Hb and HpSPHb all competed with similar IC_50_ values of 33, 40 and 41 nM, while Hb competed with a greater IC_50_ of 0.25 μM. HprSPHb showed some specific uptake at concentrations above 2 μM, which is greater than the normal serum concentration of ∼ 1 μM (*24*). Therefore, uptake experiments and binding studies were consistent, showing that all HpHb variants are equally endocytosed at low concentration, while Hp is not. Free Hb will be endocytosed at higher concentrations, which may be physiologically important in disease or infection where haemolysis results in Hp depletion.

### Calcium regulates CD163 multimer formation and HpHb uptake

The structure of the CD163 trimer bound to HpHb revealed spheres of density, which we attribute to be calcium ions, present both at the interfaces between the protomers, and at the interface between the arm of CD163_A_ and HpHb (Fig. 4a). We therefore assessed whether calcium is important for CD163 multimer formation and for HpHb binding and uptake.

**Fig. 4.**
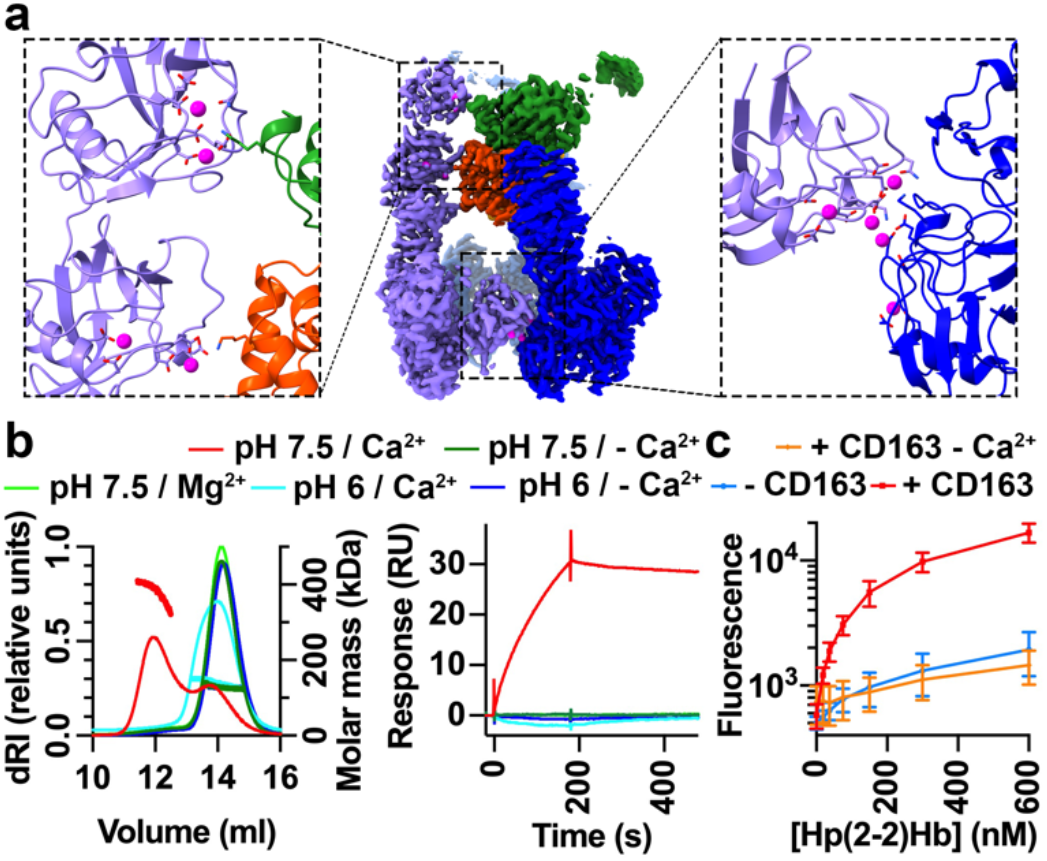
Calcium is required for CD163 multimer formation and for HpHb uptake. **a**. The central panel shows the structure of the CD163 trimer bound to HpHb, coloured as Fig. 1. The left-hand panel shows Ca2+ ions at the interface between CD163_A_ and HpHb while the right-hand panel shows Ca2+ ions at the interface between CD163_A_ and CD163_C_. **b**. The left-hand panel shows SEC-MALLS data for CD163 in the presence of Hp(1-1)Hb with the left y-axis showing the differential refractive index, while the right y-axis shows the molecular weight. This is representative of 2 repeats for measurements at pH 7.5 with Ca^2+^ and 3 repeats for all other buffers. The right-hand panel shows the binding of CD163, coated on the surface of an SPR chip, to Hp(1-1)Hb at 10 nM concentration. This is representative of 3 repeats. In both cases, the key above the graphs indicates the pH and the presence or absence of divalent cations. **c**. Measurement of the uptake of fluorescently labelled Hp(2-2)Hb into HEK293 cells expressing CD163 in the absence (red line) and presence (orange line) of EGTA. The blue line shows non-receptor-mediated uptake into cells not expressing the receptor. Each point represents the mean of three replicates and the error bars depict standard deviations.

We used SEC-MALLS to assess the multimeric state of liganded CD163. In the presence of 2.5 mM CaCl_2_ at pH 7.5, the ectodomain showed concentration-dependent multimerisation (Fig. 4b and Fig. S1c). In contrast, in the absence of calcium, at a lower pH of 6 or when CaCl_2_ is replaced by 2.5 mM MgCl_2_, CD163 was monomeric. We also used SPR to test whether the presence of calcium is required for HpHb binding. While Hp(1-1)Hb bound with an affinity of 0.28 nM in the presence of calcium (Fig. 3d), no binding was observed in the absence of calcium, in the presence of 2.5 mM MgCl_2_ or at pH 6 (Fig. 4b and Fig. S6b).

Finally, we assessed the importance of calcium for uptake into HEK293 cells expressing CD163. As the media used for these experiments contains ∼ 1.8 mM CaCl_2_, we performed the assays in the presence of 2 mM EGTA to chelate calcium. While EGTA had no effect on uptake of transferrin (Fig. S8b), suggesting that it did not negatively affect global endocytosis, it resulted in a reduction of HpHb uptake to levels observed for cells lacking CD163 (Fig. 4c). Therefore, calcium, together with pH, mediates formation of CD163 multimers and is required for HpHb uptake. As both calcium concentration and pH are low in the endosome, this suggests that CD163 forms multimers in the serum which efficiently take up ligands and that these multimers disassemble in the endosomes, facilitating HpHb release.

### Monomeric CD163 is less effective at uptake of lower-avidity ligands

Having shown that calcium is important for CD163 multimer formation, we next aimed to determine whether the putative calcium ions at the binding interface between CD163 and HpHb are also important. To achieve this, we developed a monomeric CD163 molecule. As N807 occupies a central location within the multimer interface (Fig. 5a), an R809T mutant was produced to introduce an N-linked glycosylation site at N807, sterically occluding multimer formation. This mutant remained monomeric at a concentration of 2.7 mg/ml, as measured by SEC-MALLS (Fig. 5b). SPR measurements showed that monomeric CD163 bound to Hp(1-1)Hb with an apparent affinity of 1.2 nM in the presence of 2.5 mM CaCl_2_ (Fig. 5c, Fig. S6a and Table S3), compared with 0.28 nM for the native protein (Fig. 3d). This interaction was abolished in the absence of calcium (Fig. 5c) or when CaCl_2_ was replaced with 2.5 mM MgCl_2_ (Fig. S6b). Therefore, calcium ions are important for the interaction of CD163 with its ligands.

**Fig. 5.**
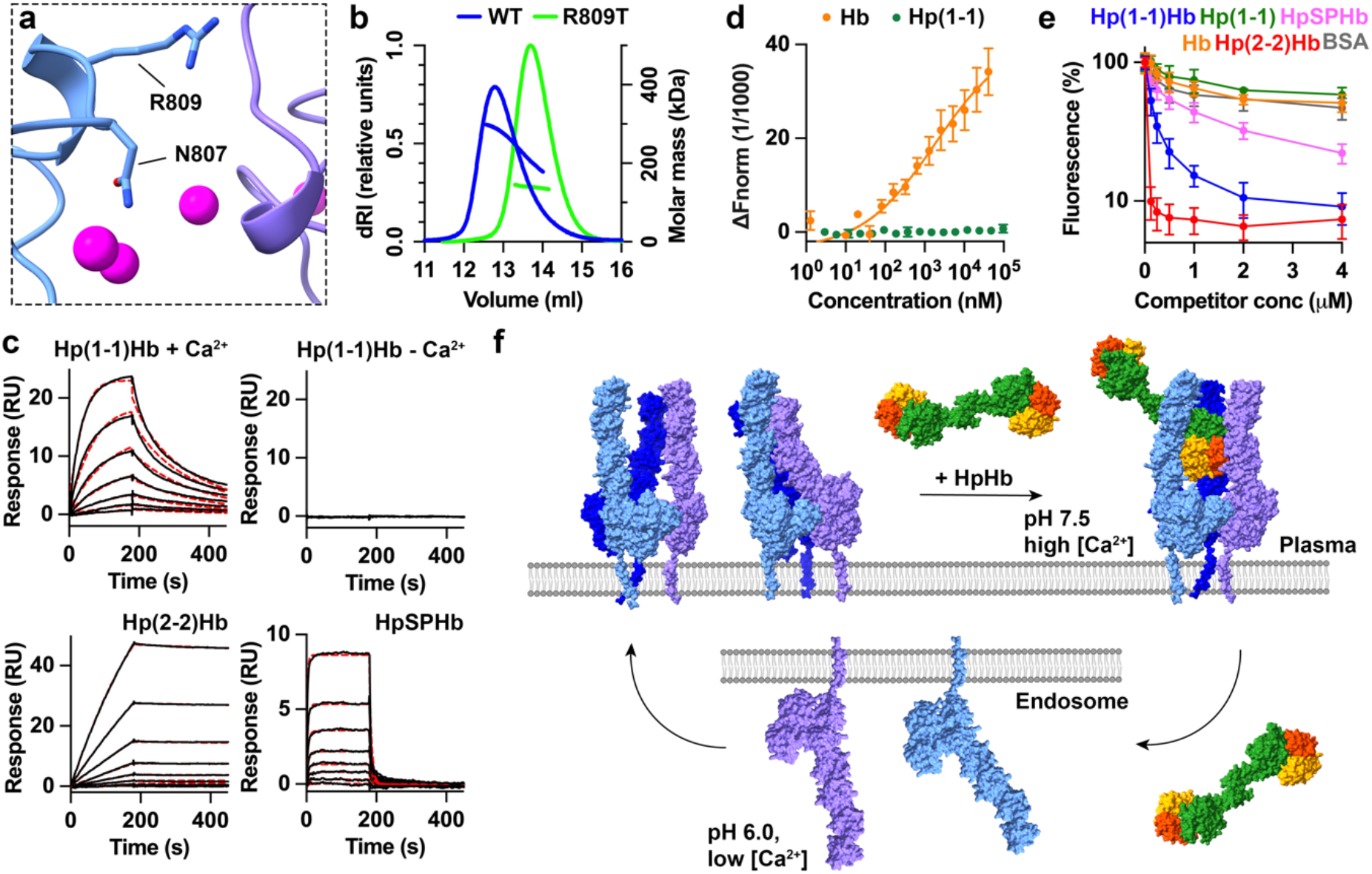
Multimerisation of CD163 allows uptake of lower-avidity ligands. **a**. A close-up of the interface between CD163_A_ and CD163_B_, showing the location of N807, which becomes modified with an N-linked glycan in the R809T mutant. **b**. SEC-MALLS analysis of wild-type (WT) CD163 ectodomain (blue) and the ectodomain of the R809T mutant (green) with the left y-axis showing the differential refractive index, while the right y-axis displays the molecular weight. This is representative of 2 repeats at a concentration of 2.7 mg/ml. **c**. Assessment of the binding of CD163-R809T ectodomain, immobilised through a C-terminal biotin, to Hp(1-1)Hb, Hp(2-2)Hb and HpSPHb by SPR analysis. In the presence of Ca^2+^, the sensorgrams represent two-fold dilution series from a maximum concentration of 5 nM for Hp(1-1)Hb, 2.5 nM for Hp(2-2)Hb and 80 nM for HpSPHb. In the absence of Ca_2+_, one injection of Hp(1-1)Hb at 10 nM is displayed. Data are shown as black lines while fitting to a one-to-one binding model is depicted as dashed red lines. These are representative of n = 3. **d**. Assessment of the binding of Hp (green) and Hb (orange) to CD163-R809T ectodomain using MST. Each point represents the mean of three replicates and the error bars are the standard deviation. **e**. Measurement of the ability of different ligands to compete for the uptake of fluorescently labelled Hp(2-2)Hb into HEK293 cells transfected with CD163-R809T. BSA is included as a control for CD163-independent effects. Each point depicts the mean of three replicates and the error bars are standard deviations. **f**. A model for the CD163-mediated uptake of HpHb. At the cell surface, calcium mediates multimerisation of CD163. Two states exist in equilibrium, with the arms either free to bind to ligand or forming arm-arm interactions which occlude ligand-binding sites. On addition of ligand, the free arms of CD163 mould around the ligand to form a complex. On internalisation into the endosome, a drop in calcium concentration and pH results in monomerisation of CD163 and ligand release. CD163 can then be recycled back to the cell surface while HpHb is degraded and detoxified.

The availability of monomeric CD163 also allowed us to assess the degree to which multimer formation is important for binding and uptake of diverse ligands. We first measured the affinities of different ligands for the R809T mutant, using SPR for HpHb complexes and MST for Hp and Hb. We found that apparent affinities for monomeric CD163 differed more than 1000-fold between Hp(2-2)Hb and HpSPHb, with values at 31 pM and 38 nM, respectively (Fig. 5c, Fig. S6a and Table S3). This stands in contrast to the similar affinities observed between HpHb variants for multimeric CD163 (Fig. 3d). The affinity for Hb was estimated to be 1.9 μM, a 10-fold decrease compared to the wild-type receptor and no Hp binding was detected at 96 μM CD163 monomer (Fig. 5d, Fig. S7 and Table S3). Therefore, monomeric CD163 bound to all ligands, other than Hp(2-2)Hb, with a lower apparent affinity than multimeric CD163.

To assess the impact of the CD163 multimer formation on the uptake of various ligands, we generated a HEK293 cell line in which full-length monomeric R809T was expressed. This allowed us to measure the internalisation of fluorescently labelled Hp(2-2)Hb and to assess how effectively it competes for the uptake of non-fluorescent ligands (Fig. 5e). Fluorescent Hp(2-2)Hb was taken up into cells expressing monomeric CD163 with a similar concentration dependence as into cells which express wild-type CD163 (Fig. S8a). However, while wild-type CD163 showed equivalent uptake of all HpHb variants, and weaker uptake of Hb (Fig. 3f), monomeric CD163 showed uptake of HpHb which varied with the number of HpSPHb heads, with most effective uptake of Hp(2-2)Hb, then Hp(1-1)Hb and finally HpSPHb, with IC_50_ values of 27 nM, 0.14 μM and 0.64 μM (Fig. 5e). Monomeric CD163 also showed decreased uptake of Hb, with an IC_50_ larger than 4 μM. Therefore while the uptake of high-avidity Hp(2-2)Hb can be mediated by monomeric CD163, multimer formation allows the efficient uptake of lower-affinity or -avidity ligands, making CD163 a more promiscuous scavenger receptor.

## Discussion

CD163 is the archetypal example of a type I scavenger receptor, with a primary known role in the detoxification of Hb released by erythrocyte damage (*12*). To achieve this, it internalises the ‘essentially irreversible’ complex (*6, 27*) between cell-free Hb and serum Hp into macrophages. While free Hb rapidly forms complexes with Hp under healthy conditions, infectious or genetic diseases which increase erythrocyte lysis are associated with Hp depletion. In these circumstances, it is beneficial for CD163 to mediate free Hb uptake. In contrast, uptake of Hp is not beneficial, as it would deplete Hp from the plasma without contributing to Hb detoxification. How does CD163 selectively recognise different HpHb isoforms, as well as Hb, without recognising Hp? The findings presented here allows us to propose a model for how CD163 mediates specific binding and release of diverse ligands (Fig. 5f).

Our structural studies of CD163 show how it mediates ligand specificity, allowing uptake of ligands with diverse stoichiometries and architectures. Two or three CD163 protomers form an assembly, with the higher local concentrations on the membrane surface most likely favouring trimers. Each protomer presents an arm which provides potential ligand-contact surfaces and these arms come together in different conformations to create a ligand-binding site for a single HpHb head. This ensures that HpHb variants with different numbers of heads are all endocytosed equally. The interaction between HpHb and CD163 involves eight SRCR domains, which mould around the ligand. This formation of a binding site by bringing together multiple small binding sites will allow promiscuity, with different combinations of these small surfaces creating binding sites for different ligands. Together with flexibility in the trimeric base, this ensures that binding sites can mould to accommodate different ligands and allows a trimeric receptor to recognise asymmetric binding partners. Most of the interactions form between CD163 and the Hb dimer with fewer interactions with Hp. This leads to uptake of free Hb but not unnecessary depletion of free Hp. The CD163 arms can also interact with one another, using the same surfaces which form ligand-binding sites. These arm-arm interactions will compete with arm-ligand interactions and it is likely that this underpins a mechanism of autoinhibition, with weaker ligands being unable to compete with arm-arm interactions for receptor binding, therefore dampening the extent of CD163 promiscuity.

The structure also reveals the role of calcium ions, which mediate two distinct functions. Calcium ions are found at the interfaces between protomers in the base of CD163, mediating multimer formation. This allows the uptake of lower-avidity ligands and scavenger receptor promiscuity. The predominance of ion-mediated interactions at these interfaces also gives the interface rotational flexibility, allowing protomers to adopt different relative conformations as the receptor moulds around a ligand. In addition, calcium ions are found within the binding sites between arms and ligands and this most likely makes each of these small interfaces more promiscuous, with stronger electrostatic binding rather than more extensive shape complementarity driving the interaction. Finally, the use of calcium in critical interfaces, together with pH, is important for ligand release. At the calcium concentrations in sera, CD163 will multimerise and bind ligands, while in low calcium, lower pH conditions in the endosomes, ligands will be released for degradation. Therefore, these studies show how the CD163 scavenger receptor is built for promiscuous ligand binding and release, revealing how it mediates the essential physiological process of Hb detoxification.

## Materials and Methods

### Expression and purification

The ectodomain of human CD163 (Uniprot Q86VB7-1 residues 46-1050) was cloned into pHLsec(*28*) and expressed with a C-terminal GAA-linker and a C-tag using the Expi293 Expression System (Gibco) according to the manufacturer’s instructions. Cell supernatant was harvested 6 days after transfection, passed through a 0.45 μm filter and applied to CaptureSelect C-tagXL Affinity Matrix (Thermo Scientific). Subsequently, the resin was washed with 30 column volumes (CV) of 20 mM Tris pH 7.5, 150 mM NaCl, and CD163 was eluted using 5 CV of 20 mM Tris pH 7.5, 2 M MgCl_2_. The protein was further purified on a Superose 6 10/300 column (Cytiva) into 20 mM HEPES pH 7.5, 150 mM NaCl, 2.5 mM CaCl_2_, flash frozen in liquid nitrogen and stored at – 80 °C. CD163-BAP was generated by fusing CD163 to a BAP tag, a GAA linker and the C-tag, and was expressed in the presence of 0.1 mM biotin. Site-directed mutagenesis of CD163-BAP was used to generate the monomeric R809T variant. Both were purified as described for CD163.

### Production of HpHb complexes

Hb was obtained by injecting the supernatant of freshly lysed human erythrocytes on a Superdex 75 16/600 column (Cytiva) which was run using 20 mM HEPES pH 7.5, 150 mM NaCl, 2.5 mM CaCl_2_. Hp(1-1) and Hp(2-2) extracted from human plasma were purchased from Sigma-Aldrich (SRP6507 and SRP6508). HpSP (Uniprot P00738-2 residues 89-347) and HprSP(*19*) (Uniprot P00739-1 residues 90-348) were cloned into pHLsec with a GAA linker and the C-tag, expressed in Expi293F cells for 6 and 5 days post-transfection, respectively, and C-tag purified as for the CD163 ectodomain. HpSP and HprSP were further purified on a Superdex 200 10/300 column (Cytiva) equilibrated in 20 mM HEPES pH 7.5, 150 mM NaCl, 2.5 mM CaCl_2_.

Complexes of various Hp phenotypes with Hb were formed by mixing Hp with a molar excess of Hb. Hp(1-1)Hb and Hp(2-2)Hb were directly injected on a Superdex 200 10/300 and a Superose 6 10/300 column, respectively, and eluted into 20 mM HEPES pH 7.5, 150 mM NaCl, 2.5 mM CaCl_2_. HpSPHb and HprSPHb were first subjected to C-tag purification before application on a Superdex 200 10/300 column.

The concentrations of HpHb complexes provided here referred to individual heads rather than entire HpHb molecules. Hb and Hp(1-1) concentrations were given for tetramers and dimer, respectively.

### Cryo-EM sample preparation and data collection

To prepare samples for cryogenic electron microscopy, CD163 was mixed with a molar excess of Hp(1-1)Hb, injected on a Superose 6 3.2/300 column (Cytiva) and eluted into HBS with calcium. The peak fractions containing CD163:Hp(1-1)Hb were pooled and concentrated to 1.3 mg/ml. Quantifoil R 1.2/1.3 Cu 300 grids were glow-discharged for 40 s at 15 mA and mounted in a Vitrobot Mark IV (Thermo Scientific) operated at 4 °C and 100 % humidity. Concentrated complexes were applied onto the grids, which were blotted for 2.5 – 4 s and plunge-frozen in liquid ethane. The grids were imaged on a Titan Krios G3 microscope (Thermo Scientific) equipped with a 300 kV emission gun, a K3 direct electron detector (Gatan) and a BioQuantum imaging filter (Gatan) with a slit width of 20 eV. EPU (Thermo Scientific) was used for automated data acquisition and was set to faster acquisition. Micrographs were collected with a defocus of −2.0 to −0.4 μm, a magnification of 58,149x, a pixel size of 0.832 Å and a dose of 41.68 e^−^/Å^2^ over a period of 2.5 s. Three grids were subjected to data acquisition, yielding 39,445 movies (8,303, 17,714 and 13,428, respectively).

Unliganded CD163 was concentrated to 1.3 mg/ml after gel filtration, and corresponding samples were prepared under the same conditions as CD163:Hp(1-1)Hb. Data collection was similar to that for the liganded complex, with the defocus set to −1.8 to −0.6 μm and a dose of 42.82 e^−^/Å^2^ over 2.6 s. 19,412 micrographs were collected from a single grid.

### Cryo-EM image processing

Micrographs from the CD163:Hp(1-1)Hb dataset were pre-processed in SIMPLE 3.0 (*20*). Following motion correction and contrast transfer function (CTF) parameters estimation, micrographs were picked using a template from a previous test data collection. After 2D classification, 1,573,382 particles were extracted and exported into CryoSPARC (*21*), where the particles were further classified and used for 3D reconstruction. The resulting volume was used to re-pick the full dataset, resulting in 15,666,525 particles with a box size of 512 pixels Fourier-cropped to 384 pixels.

Over the course of three rounds of 2D classification, classes containing poor particles were excluded, leaving 3,502,829 particles that were used for 3D reconstruction into seven classes. Classes for dimeric and trimeric CD163:Hp(1-1)Hb complexes containing most features underwent 3D classification via *ab initio* reconstruction and heterogeneous refinement, after which particles with nearly identical features were combined. 463,222 particles corresponding to the trimeric complex were re-extracted to the original pixel spacing of 0.832 Å and subjected to non-uniform refinement with optimisation of CTF parameters. This led to a volume at 3.2 Å resolution according to the gold-standard Fourier Shell Correlation (GSFSC) definition at 0.143. For the dimeric complex, 456,124 particles were re-extracted to remove the Fourier-crop and processed in a final non-uniform refinement step with per-particle defocus and CTF parameters optimisation, yielding a map at 2.8 Å resolution with the original pixel spacing of 0.832 Å.

Unliganded CD163 was processed similarly to CD163:Hp(1-1)Hb. After pre-processing and initial 2D classification in SIMPLE, 798,259 particles were exported to CryoSPARC. These particles were used to reconstruct a 3D template used to repick all micrographs. 13,772,487 particles were extracted using a box size of 512 pixels down-sampled to 384 pixels and sorted in two rounds of 2D classification. This resulted in 3,308,159 particles that were used to generate seven *ab initio* volumes. A high-resolution dimer class, a trimer class with disordered arms and a trimer class with arm-arm contacts were identified. Two rounds of 3D classification were performed on particles corresponding to the dimer class while particles corresponding to the trimer classes were 3D-classified once, after which particles with similar features were combined. 723,620 particles corresponding to the dimer class were re-extracted to remove down-sampling and subjected to non-uniform refinement, yielding a volume at 3.1 Å resolution with a pixel size of 0.832 Å. 81,871 particles corresponding to the trimer class with disordered arms and 432,423 particles representing the trimer with arm-arm contacts underwent non-uniform refinement. This resulted in maps at 3.8 and 3.4 Å, respectively. All maps were sharpened by DeepEMhancer (*29*) using the default settings and were depicted using ChimeraX v1.7 (*30*).

### Model building and refinement

Trimeric CD163:Hp(1-1)Hb exhibited a higher resolution for the CD163 base while subunit A of dimeric CD163:Hp(1-1)Hb most clearly resolved an arm of CD163. Therefore, SRCR domains 5-9 of the liganded complexes were built using ChimeraX by rigid-body fitting individual AlphaFold2-predicted domains (*31*) into the corresponding densities of trimeric CD163:Hp(1-1)Hb. Domains 2-4 were docked into the densities of the liganded dimer. Falsely built regions were corrected in Coot 0.9.8.8 (*32*) before docking the corrected SRCR domain models into unmodelled densities of the liganded CD163 maps. These models were then manually readjusted in Coot.

A crystal structure of HpSPHb(*19*) (PDB: 4×0L) and Hp(1-1)Hb from a crystal structure of its complex with the *T. brucei* HpHb receptor (PDB: 4WJG) were docked into ligand densities of trimeric and dimeric CD163:Hp(1-1)Hb, respectively. Both models were initially subjected to optimisation using ISOLDE (*33*) and subsequent iterative real-space refinement in PHENIX (*34*) and manual correction in Coot.

To build the unliganded CD163 dimer, SRCR domains from the liganded dimer were individually rigid-body fitted into corresponding densities using ChimeraX (*30*) and manually adjusted in Coot. The unliganded dimer was refined as the liganded complexes. PDBePISA (*35*) was used to calculate ligand-binding interfaces and interfaces of arm-arm contact. Models were displayed using ChimeraX.

### Surface plasmon resonance

SPR was performed on a Biacore T200 SPR system (Cytiva) operated at 25 °C using the Biotin CAPture Kit, Series S (Cytiva). SPR buffer was either HBST (20 mM HEPES pH 7.5, 150 mM NaCl, 0.005 % Tween-20) or MBST (20 mM MES pH 6.0, 150 mM NaCl, 0.005 % Tween-20) supplemented with either 2.5 mM CaCl_2_ or 2.5 mM MgCl_2_ or neither. Ligand and analyte were buffer exchanged using 0.5 ml Zeba Spin Desalting columns (7K MWCO, Thermo Scientific) into SPR buffer prior to experimentation.

Biotinylated CD163-WT-BAP or CD163-R809T-BAP diluted to 8 μg/ml were captured at 8 μl/min for 160 s. Under binding conditions for CD163-WT-BAP, two-fold dilution series of 10-0.078 nM Hp(1-1)Hb, 10-0.078 nM Hp(2-2)Hb, 20-0.16 nM HpSPHb or 20 nM HprSPHb were flowed through the chip at 30 μl/min for 3 min, followed by dissociation for 5 min. For CD163-R809T-BAP, two-fold dilution series of 5-0.078 nM Hp(1-1)Hb, 2.5-0.019 nM Hp(2-2)Hb and 80-0.63 nM HpSPHb were used. Under non-binding conditions, 10 nM Hp(1-1)Hb and 20 nM HprSPHb were injected onto the chip. The chip surface was regenerated by injecting 6 M guanidine hydrochloride, 0.25 M NaOH at 10 μl/min for 80 s. Under binding conditions, experiments were conducted three times with three independent dilution series. Under non-binding conditions, a single concentration corresponding to the highest concentration observed under binding conditions was injected three times. Kinetic analysis was performed using BIAevaluation software v1.0 (Cytiva). Curves were depicted using GraphPad Prism v10.3.1.

### Fluorescent labelling of CD163 ligands

For MST, Hb and Hp(1-1) were fluorescently labelled using Alexa Fluor 647 NHS Ester (Thermo Scientific) according to the manufacturer’s instructions. For uptake experiments, Hp(2-2) was labelled using Alexa Fluor 594 NHS Ester (Thermo Scientific). Unbound dye was removed by applying the sample on a Zeba Spin Desalting column and a Superdex 200 10/300 column and eluting it in HBS with calcium. Fluorescent Hp(1-1)Hb and Hp(2-2)Hb were produced by mixing fluorescent Hp(1-1) and Hp(2-2), respectively, with an excess of Hb and subsequent gel filtration.

### Microscale thermophoresis

MST was carried out on a Monolith NT.115 (NanoTemper) operated at 25 °C. Four different sets of binding partners were prepared in 20 mM HEPES pH 7.5, 150 mM NaCl, 2.5 mM CaCl_2_, incubated for an hour at room temperature and loaded into Monolith Capillaries (NanoTemper) for measurement. In the first set, 200 nM fluorescent Hb was mixed with a two-fold dilution series of CD163-WT (24.4 μM – 0.74 nM), which was measured at 10 % LED and 20 % MST power. In the second set, 500 nM Hp(1-1) was incubated with CD163-WT (37.9 μM – 1.2 nM) and was recorded at 8 % LED and 20 % MST power. In the third set, 200 nM Hb was mixed with CD163-R809T-BAP at 41.5 μM – 1.3 nM and was measured at 10 % LED and 20 % MST power. In the fourth set, 1.0 μM Hp(1-1) was mixed with 96.3 μM – 2.9 nM CD163-R809T-BAP and was analysed at 6 % LED and 20 % MST power. Every interaction was characterised in technical triplicates. Raw data was extracted from NT Analysis v1.5 (NanoTemper) and a fluorescence baseline offset was applied to individual replicates before they were combined and fitted using the ‘Sigmoidal, 4PL, X is concentration’ model in GraphPad Prism.

### Size-exclusion chromatography coupled with multi-angle laser light scattering

SEC-MALLS was conducted at room temperature on a Shimadzu Prominence UFLC System connected to a Superose 6 10/300 column, a Dawn Helios 8^+^ light scattering detector (Wyatt Technology) and an Optilab T-rEX refractive index detector (Wyatt Technology). To test the influence of the CD163 concentration on its multimeric state, duplicates of 100 μl samples containing 0.38, 0.67, 1.3 or 2.7 mg/ml CD163-WT were measured in HBS (20 mM HEPES pH 7.5, 150 mM NaCl) with 2.5 mM calcium. Similarly, duplicates with 0.25, 0.38, 0.51 or 0.76 mg/ml CD163-WT and 0.08 mg/ml Hp(1-1)Hb were analysed. To probe the role of calcium and pH, triplicates containing 0.38 mg/ml CD163-WT and 0.08 mg/ml Hp(1-1)Hb were analysed in HBS, HBS with 2.5 mM MgCl_2_, MBS (20 mM MES pH 6.0, 150 mM NaCl) or MBS with 2.5 mM CaCl_2_. To characterise the effect of R809T on the multimeric state of CD163, duplicates containing 0.38 or 2.7 mg/ml CD163-R809T-BAP were measured in HBS with calcium. Data were analysed using ASTRA v6.1 (Wyatt Technology) and plotted in GraphPad Prism.

### Generation of stably transfected, CD163-expressing HEK293 cells

The Flp-In-293 system (Invitrogen) was used to generate stably transfected, CD163-expressing cells according to the manufacturer’s instructions. The secretion signal sequence of pHLsec, full-length human CD163 (Uniprot Q86VB7-1 residues 46-1156, wild-type or R809T), a GSG linker and GFP (Uniprot P42212 residues 2-238) were cloned into pcDNA 5/FRT (Invitrogen). This expression vector and pOG44 were used to transfect ∼600,000 Flp-In-293 T-REx cells (Thermo Scientific) at a 1:9 ratio using TransIT-LT1 Transfection Reagent (Mirus Bio) according to the manufacturer’s instructions. Cells were passaged in Dulbecco’s Modified Eagle Medium (DMEM, Gibco) with 10 % fetal calf serum (FCS, Gibco) a day after transfection and selected in DMEM with 10 % FCS, 15 μg/ml blasticidin S (Gibco) and 100 μg/ml hygromycin B (VWR) two days post-transfection. After selection, cells were sorted using FACSAria Fusion (BD Biosciences) to obtain populations with high, consistent CD163 expression.

### Ligand uptake assay

For Hp(2-2)Hb uptake in the absence of competitors, ∼200,000 CD163-WT- or CD163-R809T-expressing cells or untransfected cells were washed in PBS. After incubation in DMEM with 25 mM HEPES, 0 – 600 nM Alexa Fluor 594-labelled Hp(2-2)Hb and optionally 2 mM EGTA for 30 min at 37 °C, 5 % CO_2_, cells were washed in PBS, trypsinised and washed again. Following addition of DRAQ7 (Abcam) to 3 μM, cells were measured on a LSRFortessa X-20 Cell Analyzer (BD Biosciences), equipped with a 488 and 561 nm laser to detect CD163-GFP and labelled Hp(2-2)Hb, respectively. For uptake of Alexa Fluor 594-labelled transferrin (Thermo Scientific, T13343), untransfected cells were handled as described for Hp(2-2)Hb uptake.

For uptake in the presence of competitors, CD163-WT- or CD163-R809T-expressing cells were incubated with 50 nM Alexa Fluor 594-labelled Hp(2-2)Hb and 0 – 4 μM unlabelled ligand and were processed as described above.

All uptake experiments were performed in triplicate, with cells for each sample incubated with fresh ligand and measured on three different days. To quantify uptake, the mean fluorescence of all live, single cells was calculated using FlowJo v10.10 and plotted in GraphPad Prism. In the uptake series performed with competitors, slight batch-to-batch differences were observed in the absolute fluorescence signal for the sample at 0 nM competitor. Therefore, fluorescence values in these series were normalised by defining the fluorescence at 0 nM competitor as 100 %. In competition experiments, IC_50_ values were defined as the concentration at which the fluorescence reaches 50 %.

## Acknowledgements

Cryo-EM data was collected at the COSMIC facility and we thank Rishi Matadeen, Teige Matthews-Parmer, Joe Caesar and Ed Lowe for support with data collection and processing. We acknowledge Hannah Ivison for lab management. We are grateful to David Staunton for help with biophysical methods and to Robert Hedley and Vasiliki Tsioligka for assistance with flow sorting. We thank Ian Gibbs-Seymour for advice in generation of stable cell lines and Thomas Carroll for advice on uptake experiments. We thank Dara Thaker for technical assistance.

## Data and materials availability

Cryo-EM maps are deposited in the Electron Microscopy Data Bank under accession codes EMD-52078 for dimeric CD163:HpHb, EMD-52079 for trimeric CD163:HpHb, EMD-52080 for unliganded dimeric CD163 with arm-arm contacts, EMD-52081 for unliganded trimeric CD163 with arm-arm contacts and EMD-52082 for unliganded trimeric CD163 with disordered arms. Atomic coordinates are deposited in the Protein Data Bank under accession codes 9HEJ for the liganded CD163 dimer, 9HEK for liganded trimer and 9HEL for unliganded dimer. All materials are available from the authors on request.

## Supplementary Figures

**Fig. S1.**
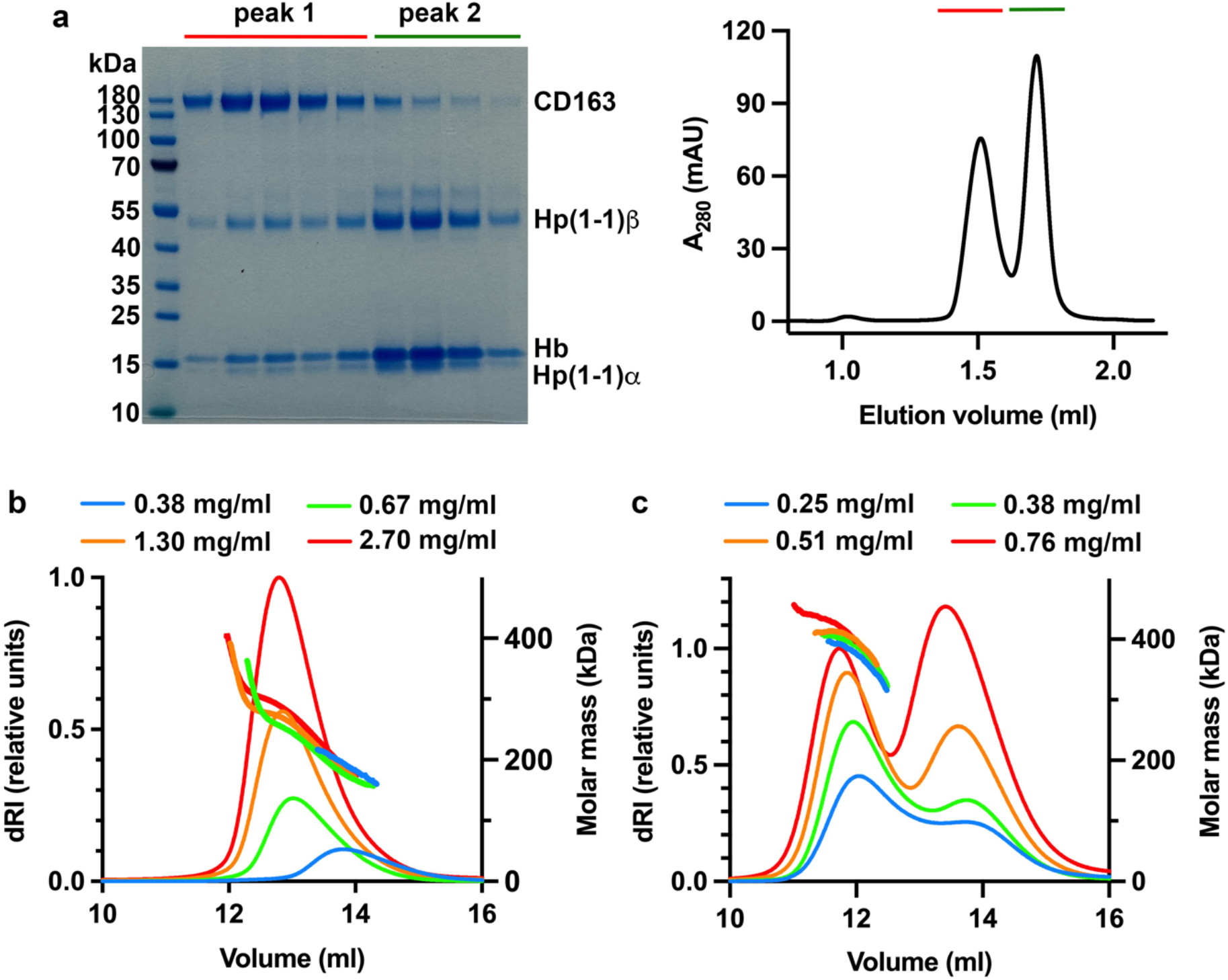
Purification and characterisation of the CD163:HpHb complex. **a**. Purification of a complex of CD163 and Hp(1-1)Hb. The right-hand panel shows the size-exclusion profile on a Superose 6 3.2/300 column while the left-hand panel shows a Coomassie-stained gel for the fractions, with peak 1 containing the liganded CD163 complex. **b**. SEC-MALLS data for unliganded CD163 at four different concentrations. The left y-axis shows the differential refractive index, while the right y-axis depicts the molecular weight. This is representative of 2 repeats. **c**. SEC-MALLS data for a complex of CD163 and Hp(1-1)Hb at four different concentrations. This is representative of 2 repeats.

**Fig. S2.**
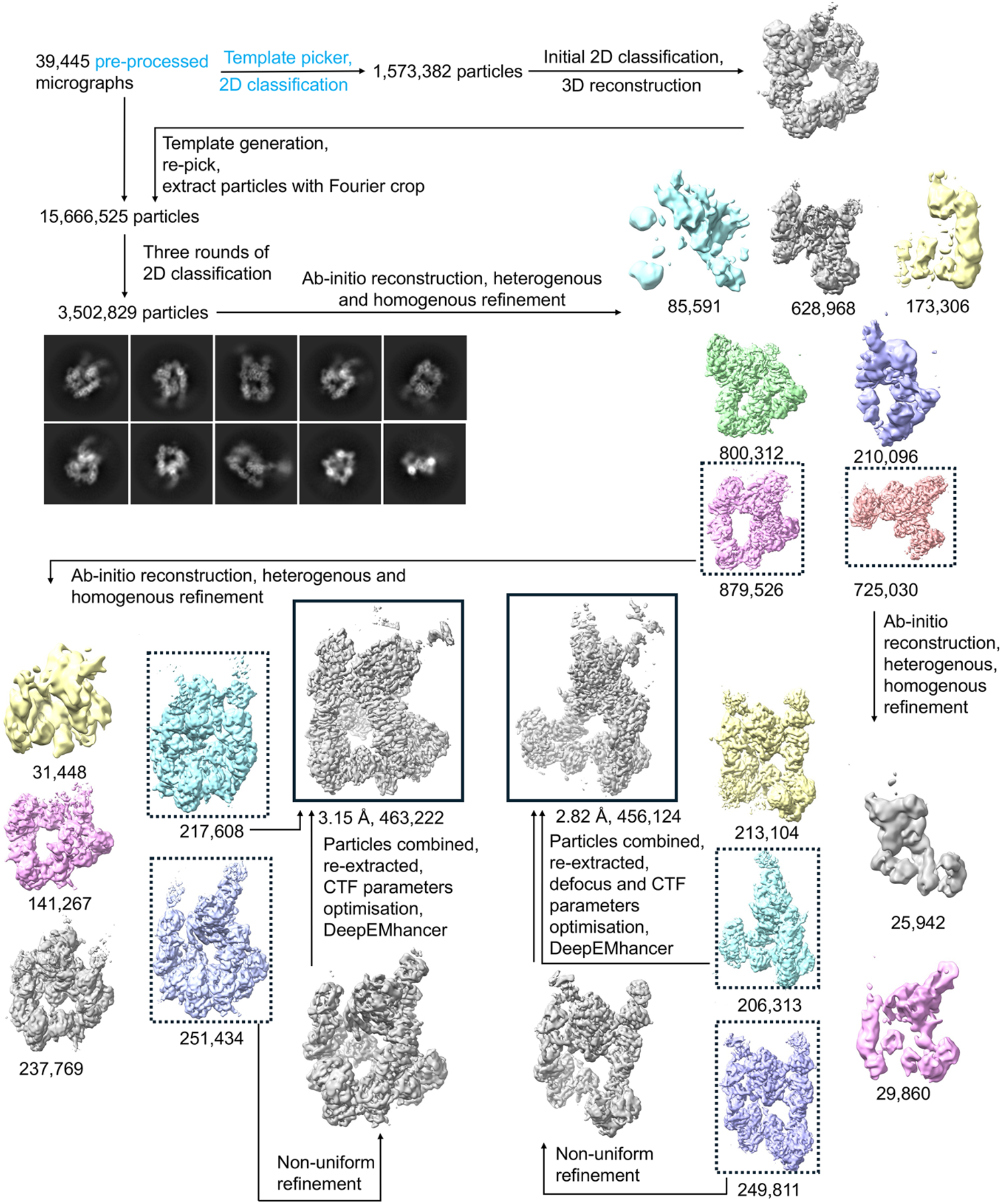
Cryo-EM image processing scheme for HpHb-bound CD163.

**Fig. S3.**
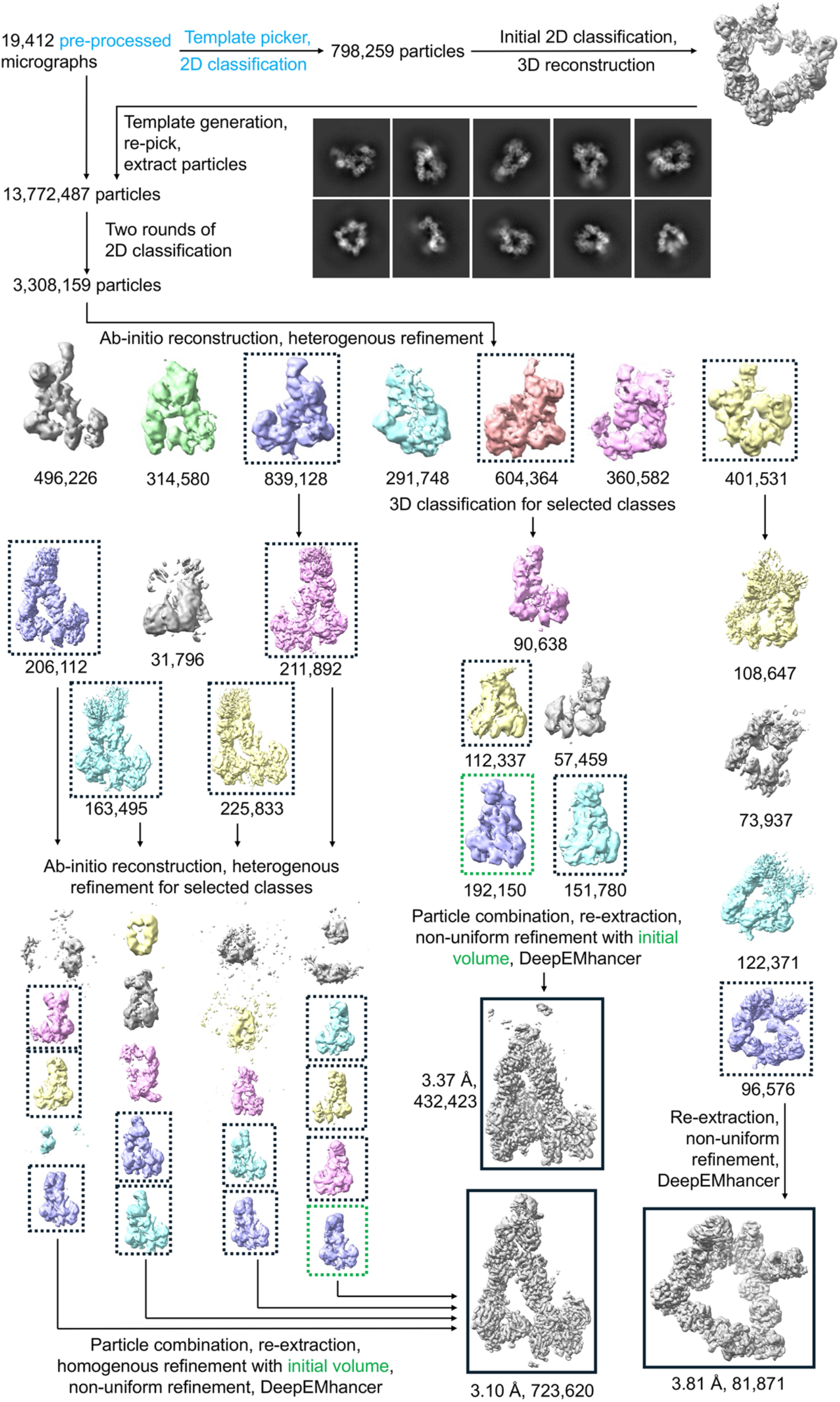
Cryo-EM image processing scheme for unliganded CD163.

**Fig. S4.**
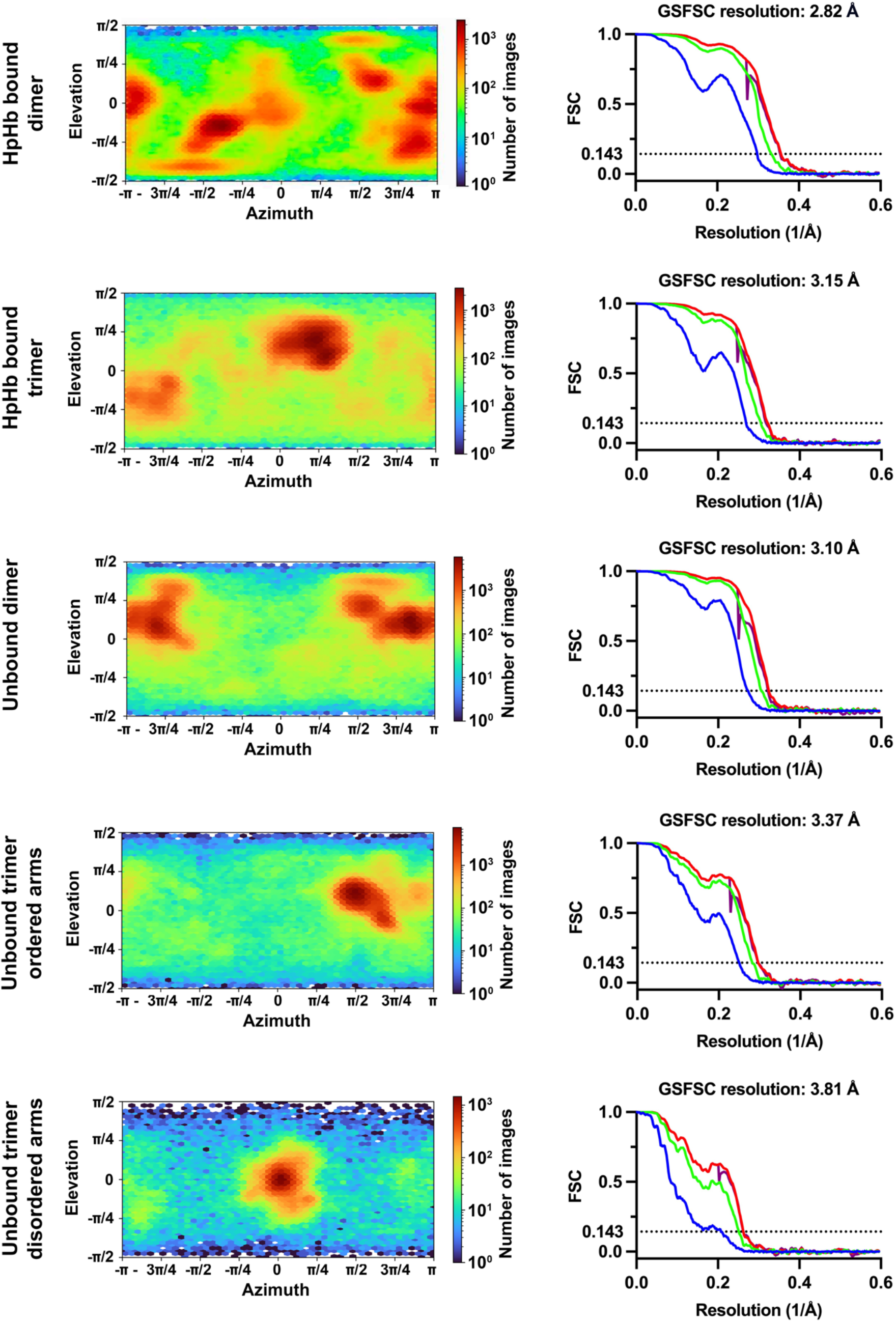
Particle-view distributions and resolutions of reconstructions. The left-hand panels show particle-view distributions for each of the five structures, while the right-hand panels display GSFSC curves.

**Fig. S5.**
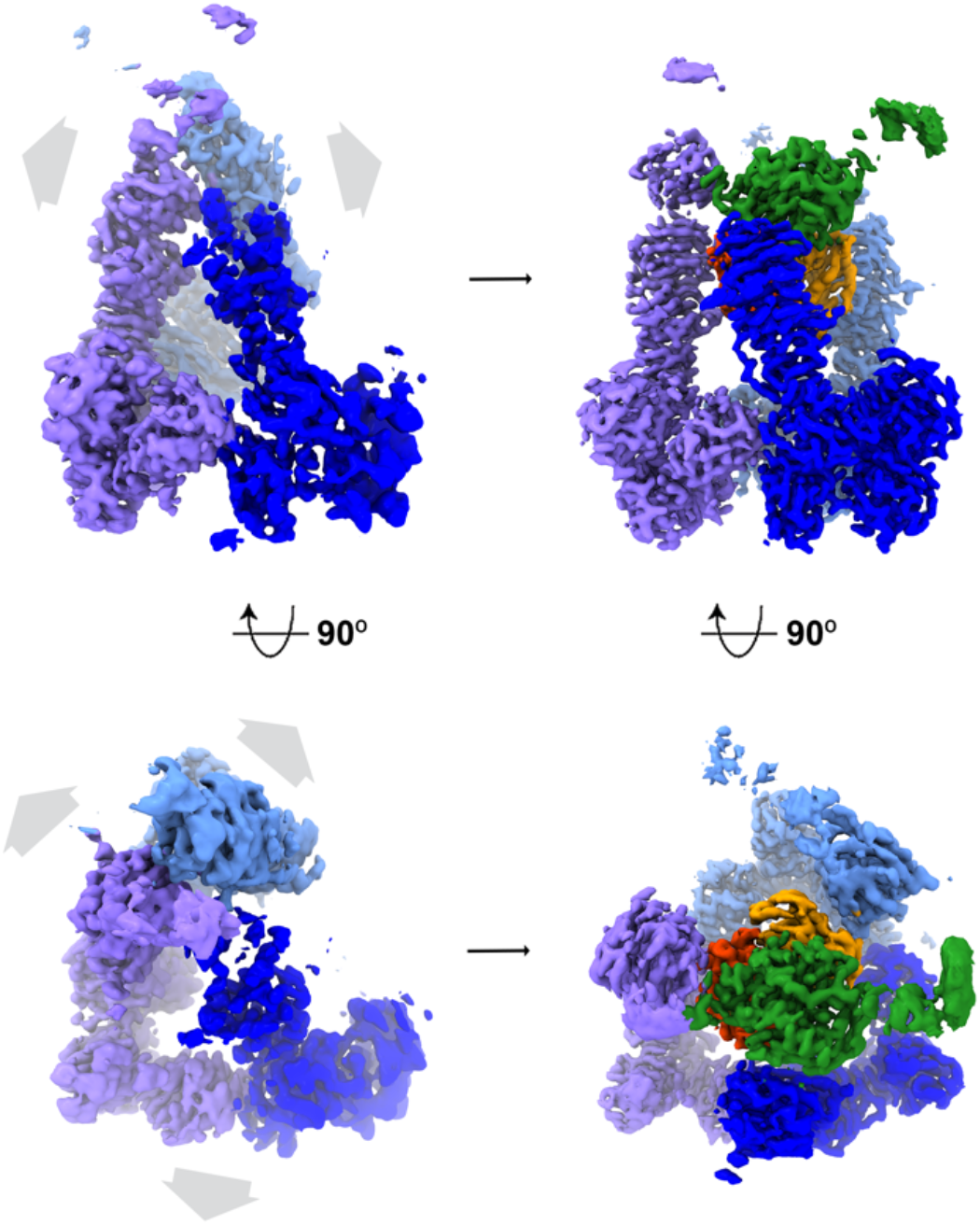
A trimeric unliganded CD163 with arm-arm interactions. The structure of a trimer of CD163 in the absence of ligand (left) and in the presence of HpHb (right), showing arm-arm contacts between the CD163 arms in the unliganded trimer, with the arms moving outwards to allow ligand binding.

**Fig. S6.**
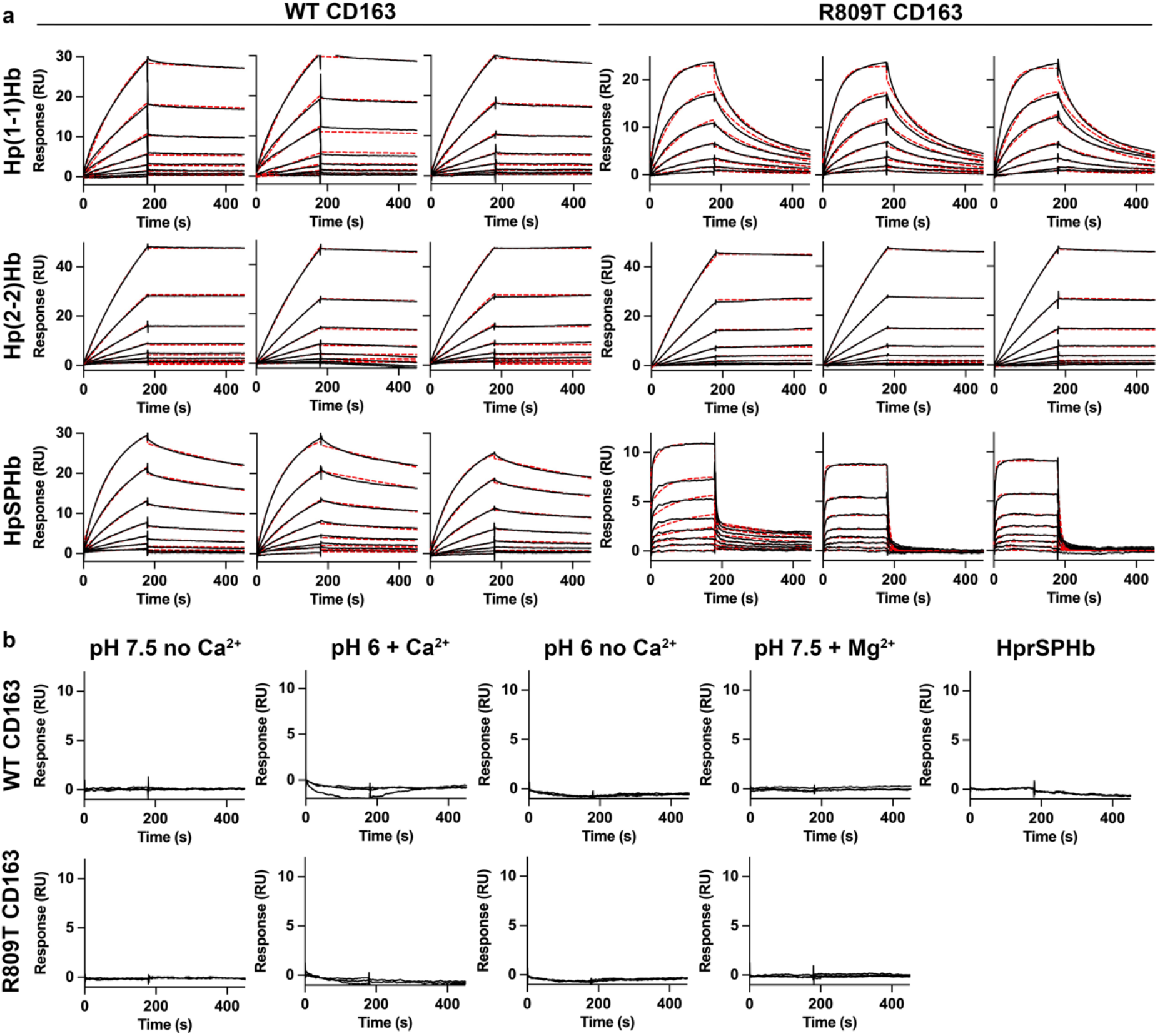
SPR analysis of ligand binding to immobilised CD163 ectodomain. **a**. SPR measurements of ligand binding. When WT CD163 was immobilised, two-fold dilution series were conducted from top concentrations of 10 nM for Hp(1-1)Hb, 10 nM for Hp(2-2)Hb and 20 nM for HpSPHb. When R809T CD163 was captured, two-fold dilution series from 5 nM Hp(1-1)Hb, 2.5 nM Hp(2-2)Hb and 80 nM HpSPHb were flowed over the chip surface. The graphs shown are the three technical replicates. **b**. SPR results in the absence of ligand binding. Three replicates are shown on every graph. The right-hand graph shows injections of HprSPHb at a concentration of 20 nM on a surface coated with WT CD163. The remaining curves show the effect of 10 nM Hp(1-1)Hb flowed over a surface coated with WT or R809T CD163, in one of four buffers: 20 mM HEPES pH 7.5, 150 mM NaCl (pH 7.5 no Ca2+); 20 mM MES pH 6.0, 150 mM NaCl, 2.5 mM CaCl_2_ (pH 6 + Ca2+); 20 mM MES pH 6.0, 150 mM NaCl (pH 6 no Ca2+) or 20 mM HEPES pH 7.5, 150 mM NaCl, 2.5 mM MgCl_2_ (pH 7.5 + Mg^2+^).

**Fig. S7.**
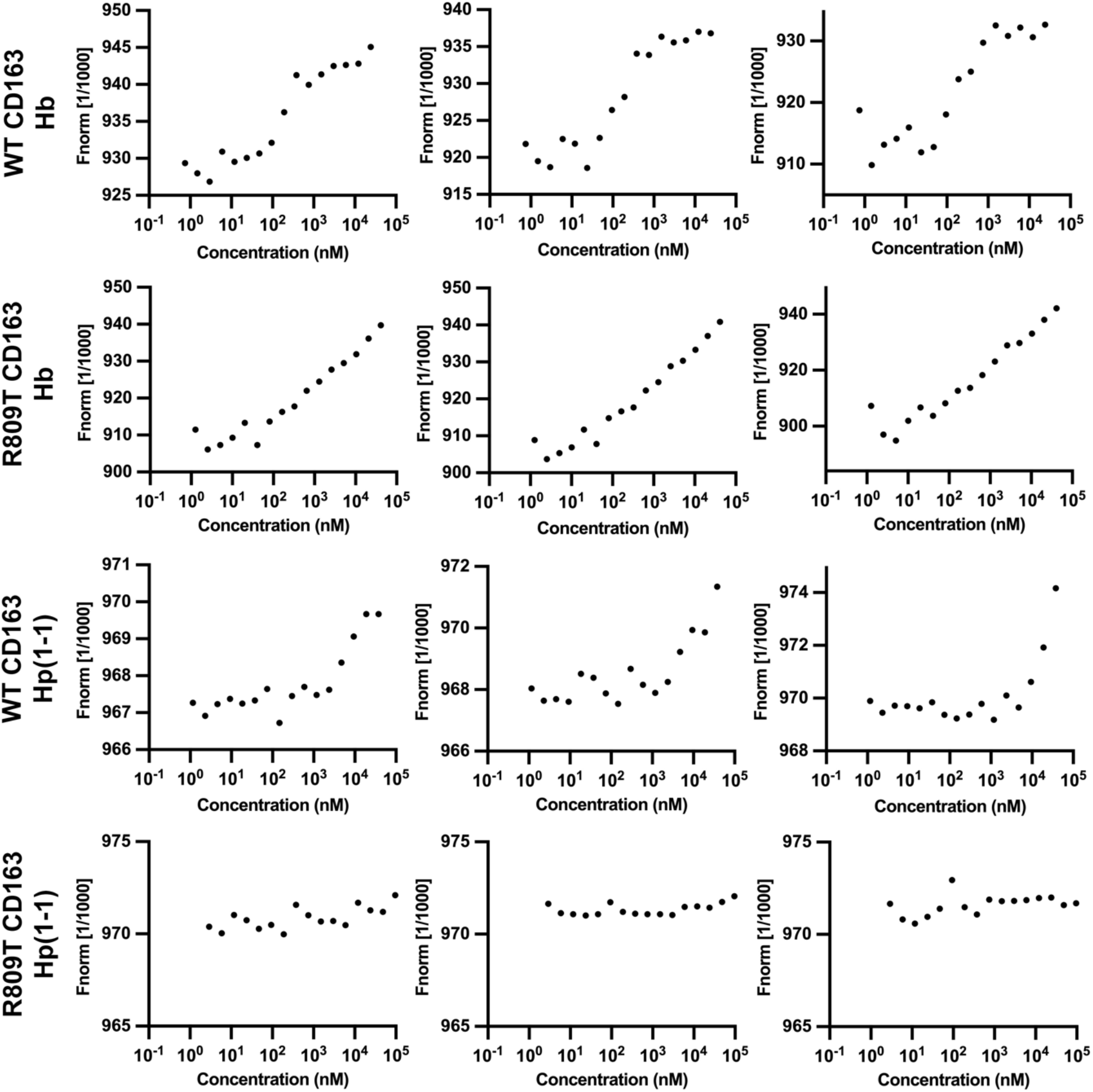
Microscale thermophoresis. MST analysis of the binding of Hp and Hb to WT CD163 or to the R809T mutant. To assess the interactions with Hb, 200 nM fluorescent Hb was mixed with two-fold dilution series of WT or R809T CD163 starting from 24.4 μM and 41.5 μM, respectively. To probe binding to Hp, 0.50 and 1.0 μM fluorescent Hp(1-1) were incubated with WT and R809T CD163 from 37.9 and 96.3 μM, respectively. Samples were measured in technical triplicates.

**Fig. S8.**
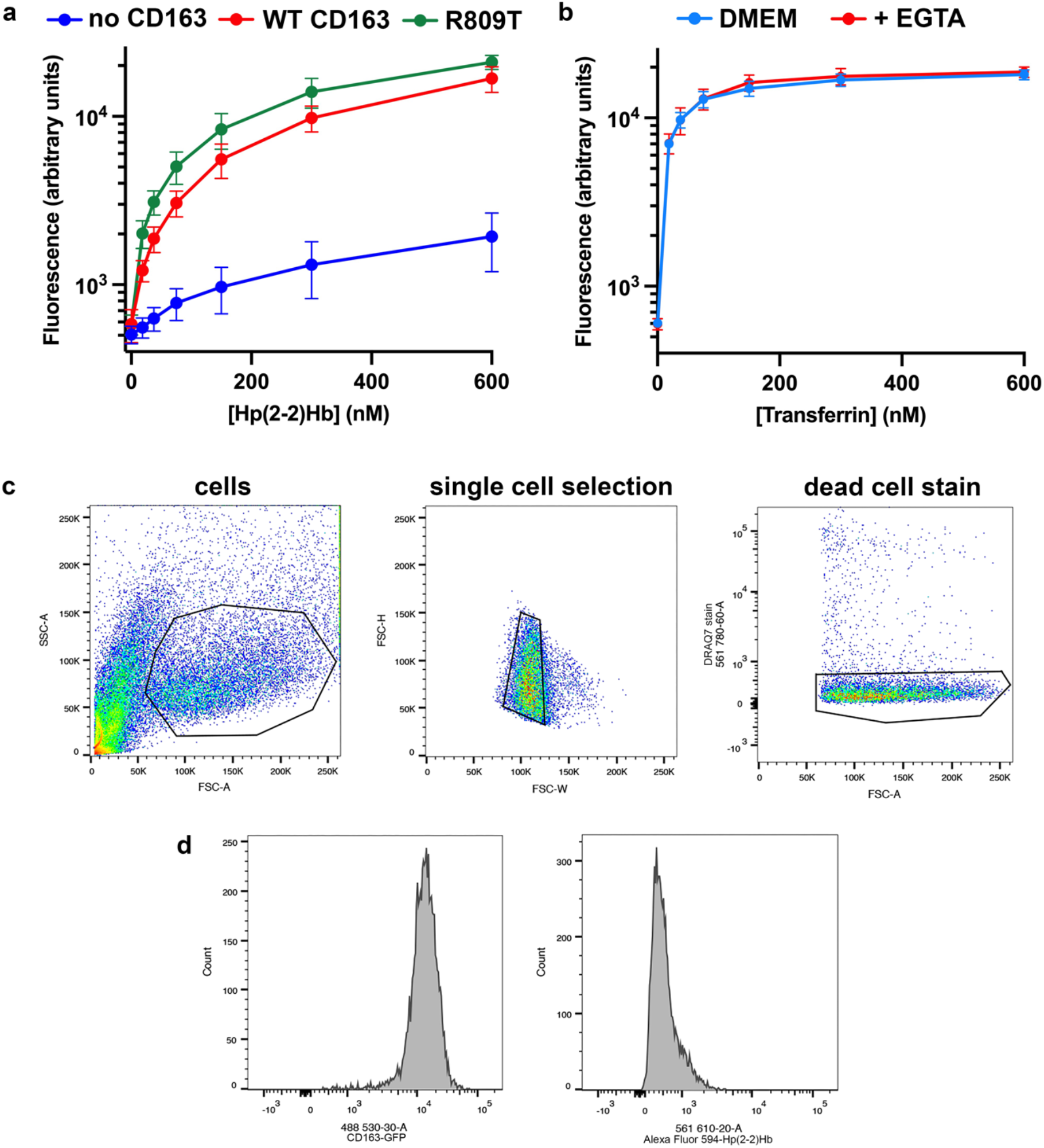
Ligand uptake analysis using fluorescence-activated cell sorting. **a**. Measurement of the uptake of fluorescently labelled Hp(2-2)Hb into untransfected HEK293 cells (blue) or into cells transfected with wild-type CD163 (red) or the R809T mutant (green). Each point represents the mean of three replicates and the error bars depict standard deviations. **b**. Measurement of the uptake of fluorescently labelled transferrin into HEK293 cells in standard DMEM media, or in DMEM with the addition of 2 mM EGTA to chelate all calcium ions. Each point represents the mean of three replicates and the error bars depict standard deviations. **c**. Gating scheme for the selection of live, single cells. The depicted gates were used for every sample in this study. **d**. Example of FACS histograms for HEK293 cells transfected with WT CD163 fused to GFP in the absence of ligand uptake. The gating strategy shown in c. was applied to obtain these histograms. CD163 expression was monitored using the left-hand histogram while ligand uptake was quantified by measuring the mean population fluorescence in the right-hand histogram.

**Table S1.** Cryo-EM data collection, refinement and validation statistics for Hp(1-1)Hb-bound CD163.

|  | Dimeric CD163:Hp(1-1)Hb | Trimeric CD163:Hp(1-1)Hb |
| --- | --- | --- |
| **Data collection and processing** |  |  |
| Microscope |  | Titan Krios |
| Detector |  | Gatan K3 with BioQuantum Imaging Filter |
| Magnification |  | 58,149 |
| Voltage (kV) |  | 300 |
| Electron exposure (e-/Å<sup>2</sup>) |  | 41.68 |
| Defocus range (μm) |  | -0.4 to -2.0 |
| Pixel size (Å) |  | 0.832 |
| Symmetry imposed |  | C1 |
| Initial particle images (no.) |  | 15,666,525 |
| Final particle images (no.) | 456,124 | 463,222 |
| Map resolution (Å) | 2.82 | 3.15 |
| FSC threshold 0.143 |  |  |
| Map resolution range (Å) | 2.25 - 35.8 | 2.55 – 27.8 |
| FSC threshold 0.5 |  |  |
| **Refinement** |  |  |
| Initial model used |  | AlphaFold2, 4X0L, 4WJG |
| Model resolution (Å) | 3.06 | 3.22 |
| FSC threshold 0.5 |  |  |
| Model composition |  |  |
| Non-hydrogen atoms | 15177 | 22064 |
| Protein residues | 1963 | 2856 |
| Ligands | 21 | 40 |
| B factors (Å<sup>2</sup>) |  |  |
| Protein | 79.41 | 80.63 |
| Ligands | 74.31 | 100.28 |
| R.m.s deviations |  |  |
| Bond lengths (Å) | 0.004 | 0.004 |
| Bond angles (°) | 0.740 | 0.756 |
| Validation |  |  |
| MolProbity score | 1.99 | 1.68 |
| Clashscore | 7.34 | 7.55 |
| Rotamer outliers (%) | 1.92 | 0.56 |
| Ramachandran plot |  |  |
| Favoured (%) | 94.7 | 96.1 |
| Allowed (%) | 5.3 | 3.9 |
| Disallowed (%) | 0.0 | 0.0 |
| Model vs. data fit |  |  |
| CC (mask) | 0.78 | 0.78 |
| CC (box) | 0.77 | 0.77 |

**Table S2.** Cryo-EM data collection, refinement and validation statistics for unliganded CD163.

|  | Unliganded dimeric<br>CD163 | Unliganded trimer<br>with disordered arms | Unliganded trimer<br>with arm contacts |
| --- | --- | --- | --- |
| **Data collection and processing** |  |  |  |
| Microscope |  | Titan Krios |  |
| Detector |  | Gatan K3 with BioQuantum Imaging Filter |  |
| Magnification |  | 58,149 |  |
| Voltage (kV) |  | 300 |  |
| Electron exposure (e-/Å<sup>2</sup>) |  | 42.82 |  |
| Defocus range (µm) |  | -0.6 to -1.8 |  |
| Pixel size (Å) |  | 0.832 |  |
| Symmetry imposed |  | C1 |  |
| Initial particle images (no.) |  | 13,772,487 |  |
| Final particle images (no.) | 723,620 | 81,871 | 432,423 |
| Map resolution (Å) | 3.10 | 3.81 | 3.37 |
| FSC threshold 0.143 |  |  |  |
| Map resolution range (Å) | 2.57 – 40.5 | 3.08 – 60.8 | 1.78 – 51.6 |
| FSC threshold 0.5 |  |  |  |
| **Refinement** |  |  |  |
| Model resolution (Å) | 3.31 |  |  |
| FSC threshold 0.5 |  |  |  |
| Model composition |  |  |  |
| Non-hydrogen atoms | 10616 |  |  |
| Protein residues | 1382 |  |  |
| Ligands | 18 |  |  |
| B factors (Å<sup>2</sup>) |  |  |  |
| Protein | 91.71 |  |  |
| Ligands | 109.73 |  |  |
| R.m.s deviations |  |  |  |
| Bond lengths (Å) | 0.003 |  |  |
| Bond angles (°) | 0.646 |  |  |
| Validation |  |  |  |
| MolProbity score | 1.78 |  |  |
| Clashscore | 8.09 |  |  |
| Poor rotamers (%) | 0.18 |  |  |
| Ramachandran plot |  |  |  |
| Favoured (%) | 95.0 |  |  |
| Allowed (%) | 5.0 |  |  |
| Disallowed (%) |  |  |  |
| Model vs. data fit |  |  |  |
| CC (mask) | 0.78 |  |  |
| CC (box) | 0.74 |  |  |

**Table S3.**
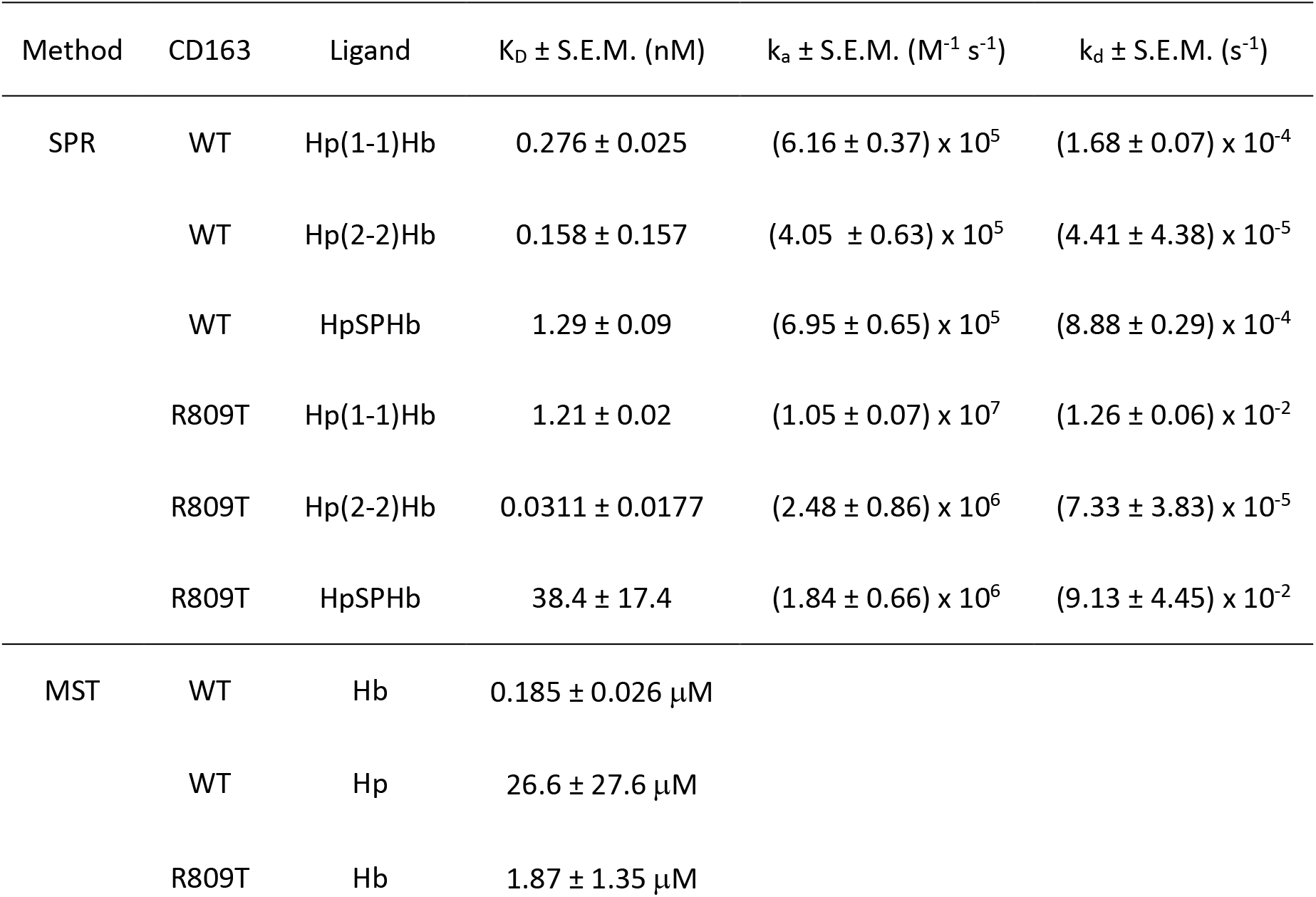
Measurement of binding constants. SPR was used to measure K_D_ and kinetic constants for the binding of WT and R809T CD163 to Hp(1-1)Hb, Hp(2-2)Hb and HpSPHb. The kinetic parameters were individually quantified using BIAevaluation (Cytiva) for every replicate, which were used to calculate the mean ± S.E.M. provided here. MST was used to determine the K_D_ values for binding of CD163 to Hb and Hp(1-1). These were derived from a fit to a curve for the combined triplicate, plotted as ΔFnorm. n = 3 for every interaction listed.

| Method | CD163 | Ligand | $K_D \pm \text{S.E.M. (nM)}$ | $k_a \pm \text{S.E.M. (M}^{-1} \text{s}^{-1})$ | $k_d \pm \text{S.E.M. (s}^{-1})$ |
| --- | --- | --- | --- | --- | --- |
| SPR | WT | Hp(1-1)Hb | $0.276 \pm 0.025$ | $(6.16 \pm 0.37) \times 10^5$ | $(1.68 \pm 0.07) \times 10^{-4}$ |
| | WT | Hp(2-2)Hb | $0.158 \pm 0.157$ | $(4.05 \pm 0.63) \times 10^5$ | $(4.41 \pm 4.38) \times 10^{-5}$ |
| | WT | HpSPHb | $1.29 \pm 0.09$ | $(6.95 \pm 0.65) \times 10^5$ | $(8.88 \pm 0.29) \times 10^{-4}$ |
| | R809T | Hp(1-1)Hb | $1.21 \pm 0.02$ | $(1.05 \pm 0.07) \times 10^7$ | $(1.26 \pm 0.06) \times 10^{-2}$ |
| | R809T | Hp(2-2)Hb | $0.0311 \pm 0.0177$ | $(2.48 \pm 0.86) \times 10^6$ | $(7.33 \pm 3.83) \times 10^{-5}$ |
| | R809T | HpSPHb | $38.4 \pm 17.4$ | $(1.84 \pm 0.66) \times 10^6$ | $(9.13 \pm 4.45) \times 10^{-2}$ |
| MST | WT | Hb | $0.185 \pm 0.026 \mu\text{M}$ | | |
| | WT | Hp | $26.6 \pm 27.6 \mu\text{M}$ | | |
| | R809T | Hb | $1.87 \pm 1.35 \mu\text{M}$ | | |

